# Amyloid beta acts synergistically as a pro-inflammatory cytokine

**DOI:** 10.1101/2020.09.28.316927

**Authors:** Thomas J. LaRocca, Alyssa N. Cavalier, Christine M. Roberts, Maddie R. Lemieux, Christopher D. Link

## Abstract

The amyloid beta (Aβ) peptide is believed to play a central role in Alzheimer’s disease (AD), the most common age-related neurodegenerative disorder. However, the natural, evolutionarily-selected functions of Aβ are incompletely understood. Here, we report that nanomolar concentrations of Aβ act synergistically with known cytokines to promote pro-inflammatory activation in primary human astrocytes (a cell type increasingly implicated in brain aging and AD). Using transcriptomics (RNA-seq), we show that Aβ can directly substitute for the complement component C1q in a cytokine cocktail previously shown to induce astrocyte immune activation. Furthermore, we show that astrocytes synergistically activated by Aβ have a transcriptional signature similar to neurotoxic “A1” astrocytes known to accumulate with age and in AD. Interestingly, we find that this biological action of Aβ at low concentrations is distinct from the transcriptome changes induced by the high/supraphysiological doses of Aβ often used in *in vitro* studies. Collectively, our results suggest an important, cytokine-like function for Aβ and a novel mechanism by which it may directly contribute to the neuroinflammation associated with brain aging and AD.

## INTRODUCTION

The amyloid beta (Aβ) peptide is centrally involved in Alzheimer’s disease (AD), the most common age-related neurodegenerative disorder [1]. However, the exact biological function and role of Aβ in the development of AD remain poorly understood. In fact, although aggregated Aβ is a well-known feature of AD pathology (i.e., in senile plaques), the peptide itself has been reported to act as a regulator of synaptic function, an antimicrobial peptide, a tumor suppressor, and a modulator of blood-brain barrier permeability [2–4]. Among these and other possible biological roles for Aβ, one compelling (and potentially unifying) idea is that Aβ may be an immune modulator. Indeed, evidence suggests that Aβ may activate pro-inflammatory signaling pathways in multiple CNS cell types [5], and this observation is consistent with the fact that neuroinflammation is a central feature of brain aging and AD [1, 6].

Neuroinflammation is characterized by increased production of pro-inflammatory cytokines in the CNS, especially by microglia and astroglia [7, 8]. It occurs in response to many stimuli (e.g., pathogens, injury, etc.) and plays an important role in protecting neurons and the brain. Growing evidence demonstrates that *chronic* neuroinflammation develops with aging [9, 10], and the neurotoxic cytokine milieu associated with this persistent neuroinflammation contributes to the development of AD [6, 11]. Innate immune activation in glial cells appears to be particularly important in this context [12, 13], and recent data demonstrate a specific accumulation of pro-inflammatory “disease-associated astrocytes” in both brain aging and AD [14]. Still, the possible mechanisms underlying astroglial activation in brain aging/AD are unclear.

One potentially simple explanation for glial cell innate immune activation in brain aging/AD could be a direct, cytokine-like effect of Aβ, which is thought to be produced largely by neurons [15, 16]. This idea has been suggested before [17], and there have been some reports of immune activation in response to Aβ [9, 18–20]. The general concept is also consistent with: 1) the fact that Aβ shares structural similarities (e.g., small amphipathic structure, expression in multiple tissues) with anti-microbial peptides like cathelicidin, which has been reported to stimulate pro-inflammatory cytokine secretion in glial cells [21]; 2) numerous studies showing that Aβ can induce pores in cell membranes [22], which is a common characteristic of immune-activating antimicrobial peptides; 3) reports that LPS (a classic innate immune stimulus) increases Aβ in the brain [23]; and 4) growing evidence that, similar to pro-inflammatory cytokines, Aβ levels increase with aging in the human brain [24, 25]. Collectively, these observations suggest that the Aβ peptide itself could have important, pro-inflammatory signaling effects in the CNS that are highly relevant to brain aging/AD. However, while aggregated Aβ can clearly induce microglial immune activation [26], the effects of soluble Aβ on human glial cells have not been extensively investigated. Here we used transcriptome analyses to investigate the global effects of soluble Aβ on primary human astrocytes. We observed that exposing these cells to 1 µM Aβ_1-42_ caused transcriptional changes indicative of immune activation. However, this response was absent with more physiological levels of Aβ (10 nM), leaving open the question of whether Aβ can truly act as a pro-inflammatory cytokine in the early stages of brain aging/AD (i.e., before higher concentrations/aggregates of the peptide develop).

Interestingly, others have shown that a cocktail of microglial-produced cytokines (tumor necrosis factor α [TNF-α], Interleukin 1α [IL-1α], and complement protein 1q [C1q]) can convert mouse astrocytes into an immune activated, neurotoxic “A1” state [27]. The same group also demonstrated that these “A1” immune-activated astrocytes (marked by expression of complement protein C3) increase with aging in mice [28] and are abundant in AD patient brains [27], suggesting that reactive astrocytes could directly contribute to age-related AD. These observations highlight the point that pro-inflammatory cytokines act in concert *in vivo* [29], which could suggest that the pro-inflammatory activity of Aβ might require other, synergistic cytokines. In support of this idea, we show here that primary human astrocytes can be converted to a reactive, A1-like state using either the originally reported TNF-α+IL-1α+C1q cocktail, or substituted cocktails in which 10 nM Aβ replaces C1q or IL-1α. Our findings suggest that immune activation may be a natural biological function of Aβ that contributes to coordinated pro-inflammatory responses in the CNS, and this could have important implications for our understanding of early mechanisms in age/AD-related neuroinflammation.

## RESULTS

### Immunomodulatory effects of Aβ on human astrocytes

In initial studies, we treated primary human astrocytes with 1 µM Aβ_1-42_ (a high but commonly used dose) and performed RNA-seq to profile changes in gene expression. We found that this high dose of Aβ differentially affected transcript levels of numerous genes. In fact, Aβ significantly increased the expression of >2600 genes and reduced >2700 genes vs. controls (**Figure 1A**, next page). Consistent with previous reports that Aβ may be pro-inflammatory, gene ontology analyses indicated that these Aβ-induced gene expression changes included increased transcription of inflammatory genes, marked by enrichment for gene ontology terms related to immune/defense responses, cytokine signaling and interferon responses (**Figure 1B**). A coherent biological response was not as obvious in the downregulated transcriptional modules, although several gene ontology terms were associated with ion transport and neurotransmitter/synapse modulation (**Figure 1B**). These findings are in line with studies that have documented the production of pro-inflammatory cytokines and reactive oxygen species (immune defense mediators) by astrocytes in response to high dose Aβ [30–32], and with more recent reports of immune/inflammatory gene expression in astrocytes isolated from AD patient brains [33]. However, when we replicated these studies using a more physiological level of Aβ_1-42_ (10nM), we observed no transcriptome/inflammatory gene expression changes (**Figure 1C**).

**Figure 1.**
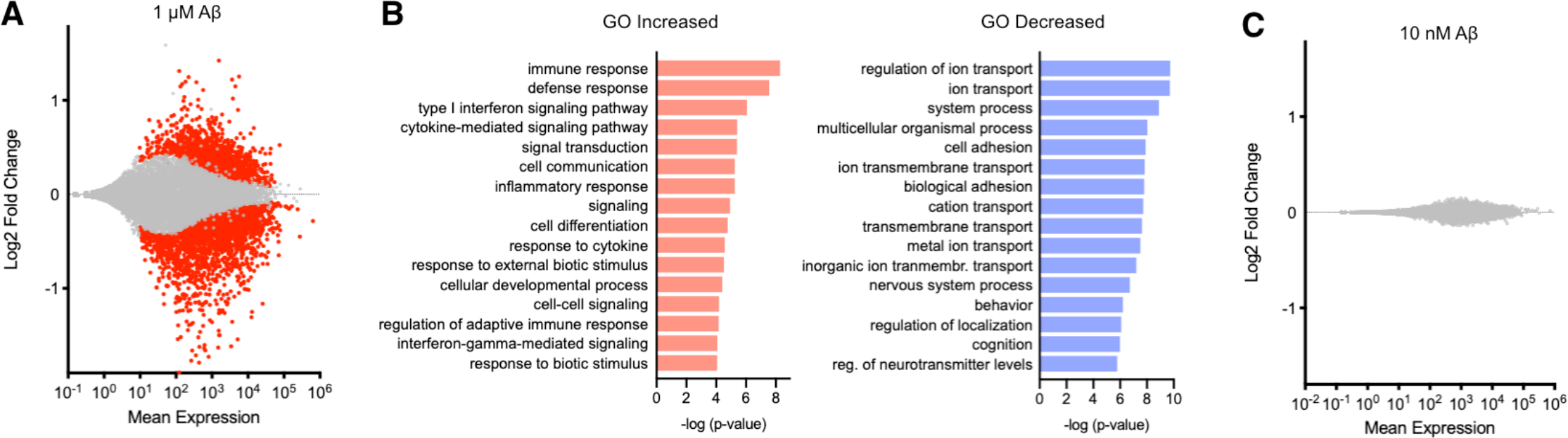
Transcriptome analysis of human astrocytes treated with high and low concentration Aβ. **(A)** MA plot showing Log2 fold change and mean gene expression levels in primary human astrocytes treated with 1 µM Aβ for 24 h. Note >5000 significantly increased/decreased genes shown in red (FDR<0.1). **(B)** Top transcriptional modules increased/decreased in gene ontology (GO) analysis of 1 µM Aβ-treated astrocytes. **(C)** MA plot showing Log2 fold change and mean gene expression (no significant changes) in astrocytes treated with 10 nM Aβ for 24 h. All experiments performed in triplicate with cells from one donor.

### Evidence for cytokine-like effects of Aβ

We considered the possibility that 10 nM Aβ_1-42_ failed to elicit immune activation in astrocytes because, *in vivo*, Aβ might act in concert with other pro-inflammatory cytokines. Building on work by Liddelow et al. [27], who demonstrated that a cocktail of three microglial-produced cytokines (TNF-α, IL-1α and C1q) was required for strong immune activation of murine astrocytes, we tested the idea that more physiological levels of Aβ might act synergistically with pro-inflammatory cytokines that are linked with aging/AD and astrocyte immune activation during aging [28]. First, we tested permutations of the TNF-α+IL-1α+C1q cocktail [27] in which we substituted Aβ for individual cytokines (all in serum-free media). Using immunofluorescence staining for CD44, a cellular adhesion molecule that tracks with astrocyte immune activation [27, 34, 35], we found that: 1) the cytokine-only TNF-α+IL-1α+C1q cocktail does, in fact, activate human astrocytes; and 2) we could substitute Aβ for C1q or IL-1α and achieve similar astrocyte activation (**Figure 2A**, next page). We then repeated these experiments and immunoblotted for additional proteins, and we found that the synergistic effect of Aβ in this cytokine cocktail was greatest when substituting it for C1q (i.e., TNF-α+IL-1α+Aβ), as indicated by greater expression of glial fibrillary acidic protein (GFAP, a common marker of reactive astrocytes) and IL-1β (a pro-inflammatory cytokine released by immune-activated glial cells) [13] (**Figure 2B**). In these exploratory trials, we used both Aβ and oligomerized Aβ (which may be more toxic [36], but that we had not tested at first); we found greater activation with non-oligomerized peptide and therefore used this in subsequent, quantitative studies, as follows: Using IL-1β and intercellular adhesion molecule 1 (ICAM-1) as indicators of AD-relevant immune activation [37–39], we found in a dose/response experiment that low concentration Aβ_1-42_ (but not as low as 1 nM) could fully substitute for C1q in activating human astrocytes (**Figure 2C**). In fact, we found that ~10 nM Aβ combined with similar concentrations of TNF-α and IL-1α was sufficient to markedly increase levels of both IL-1β and ICAM-1, both of which were not increased in response to high-dose (1 µM) Aβ (i.e., in **Figure 1**).

**Figure 2.**
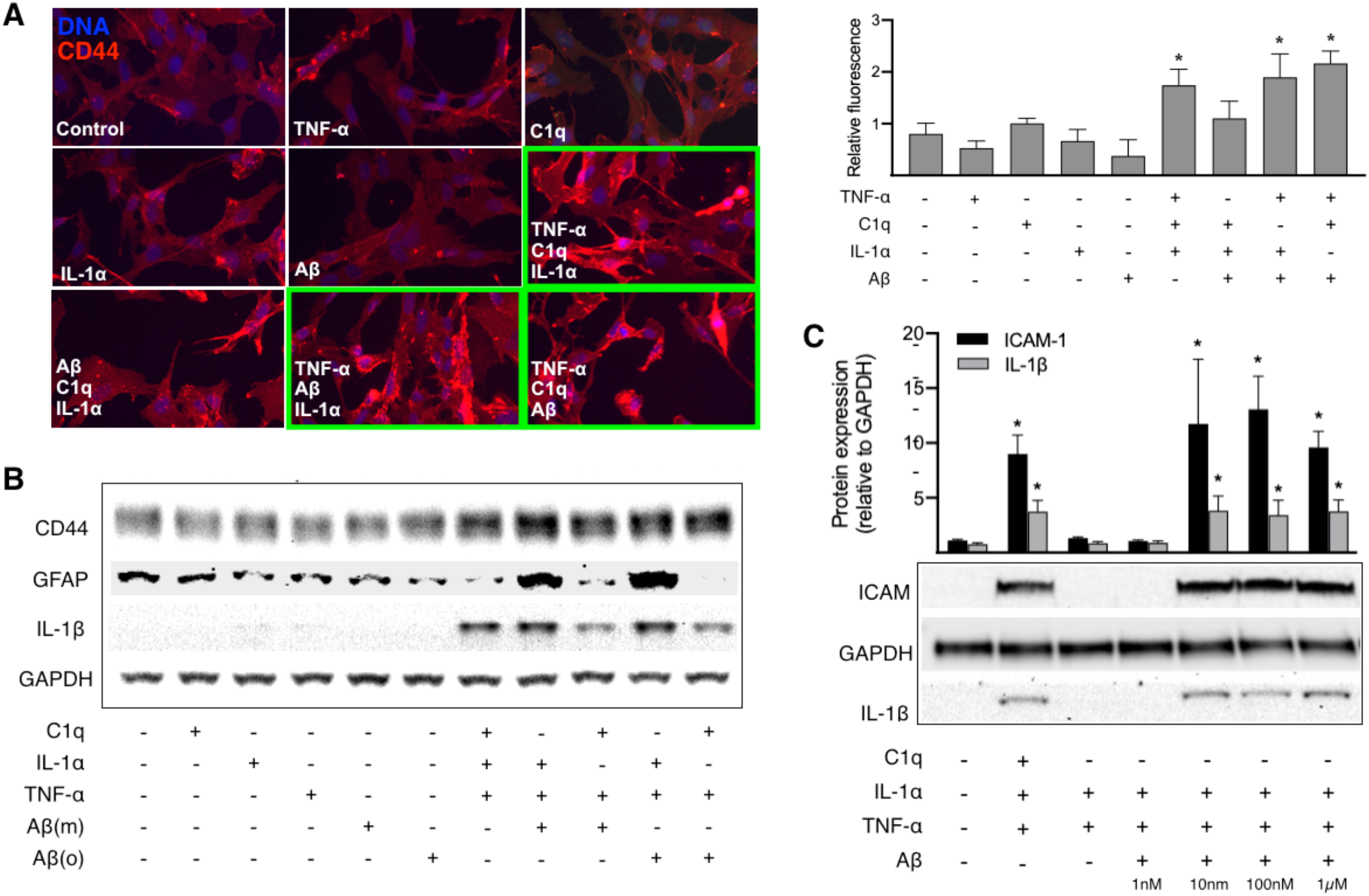
Astrocyte immune activation induced by cytokines and Aβ/cytokine cocktails. **(A)** Representative immunofluorescence staining for the astrocyte activation marker CD44 in astrocytes treated with the cytokines TNFα, IL-1α and C1q, as well as Aβ (all alone or in combination). Note greater staining with all-cytokine cocktail and when Aβ is substituted for IL-1α or C1q (relative signal quantified at right). **(B)** Western blots showing that Aβ monomers (m) and oligomers (o) can substitute for IL-1α or C1q to induce immune activation, marked by increased expression of the pro-inflammatory cytokine IL-1β and the astrocyte activation marker GFAP. Note somewhat greater activation when Aβ is substituted for C1q. **(C)** Dose-response experiments showing that nanomolar concentrations of Aβ are sufficient to cause astrocyte activation (increased expression of IL-1β and the reactive astrocyte marker ICAM-1) when combined with TNF-α and IL-1α. All experiments performed 3-5 times using primary human astrocytes from one donor. *P<0.05 vs. control, unpaired two-tailed t-test.

### Aβ/cytokine-induced transcriptome changes

To extend our analysis of human astrocyte activation beyond the limited set of markers employed in Figure 2, we repeated our experiments using RNA-seq to capture the global effects of Aβ/cytokine exposure. In these studies, we also used astrocytes from three different donors to identify the most conserved effects of the treatment. As we previously observed in replicate cultures from a single donor (**Figure 1C**), we found that 10 nM Aβ_1-42_ alone had minimal effects on transcript accumulation (**Figure 3A**, next page). Similarly, and in agreement with previous reports [27], TNF-α combined only with IL-1α also caused minimal transcriptome changes (**Figure 3B**). However, the addition of either C1q or Aβ to this cocktail resulted in a major biological response, reflected by significantly increased/decreased expression of ~2500 genes (**Figure 3C**,**D**). Gene expression patterns induced by the cytokine-only cocktail (TNF-α+IL-1α+C1q) and the Aβ/cytokine cocktail (TNF-α+IL-1α+Aβ) were strikingly similar, as reflected by no major differences (**Figure 3E**) and strong inter-treatment correlation (**Figure 3F**), again suggesting that Aβ could have cytokine-like effects. Moreover, as expected from the highly similar transcriptome patterns induced by these two treatments, gene ontology analysis revealed very similar gene set enrichments (**Figure 3G**). In particular, both treatments upregulated transcriptional modules related to immune system activation, cytokine/interferon signaling and inflammatory responses, whereas common downregulated transcriptional modules included gene sets modulating neurogenesis, synaptic structure, and other cellular health/development pathways. Interestingly, gene expression changes induced by the Aβ/cytokine cocktail only correlated weakly with those induced by high dose (1 µM) Aβ alone (**Figure 3H**). Many of the genes most affected by the Aβ/cytokine cocktail but not by Aβ alone were mediators of innate immune responses (e.g., C3 and IL-1β) and/or established markers of reactive astrocytes (e.g., lipocalin-2, LCN2) [27, 40], further suggesting that physiological Aβ may have a distinct role in immune activation.

**Figure 3.**
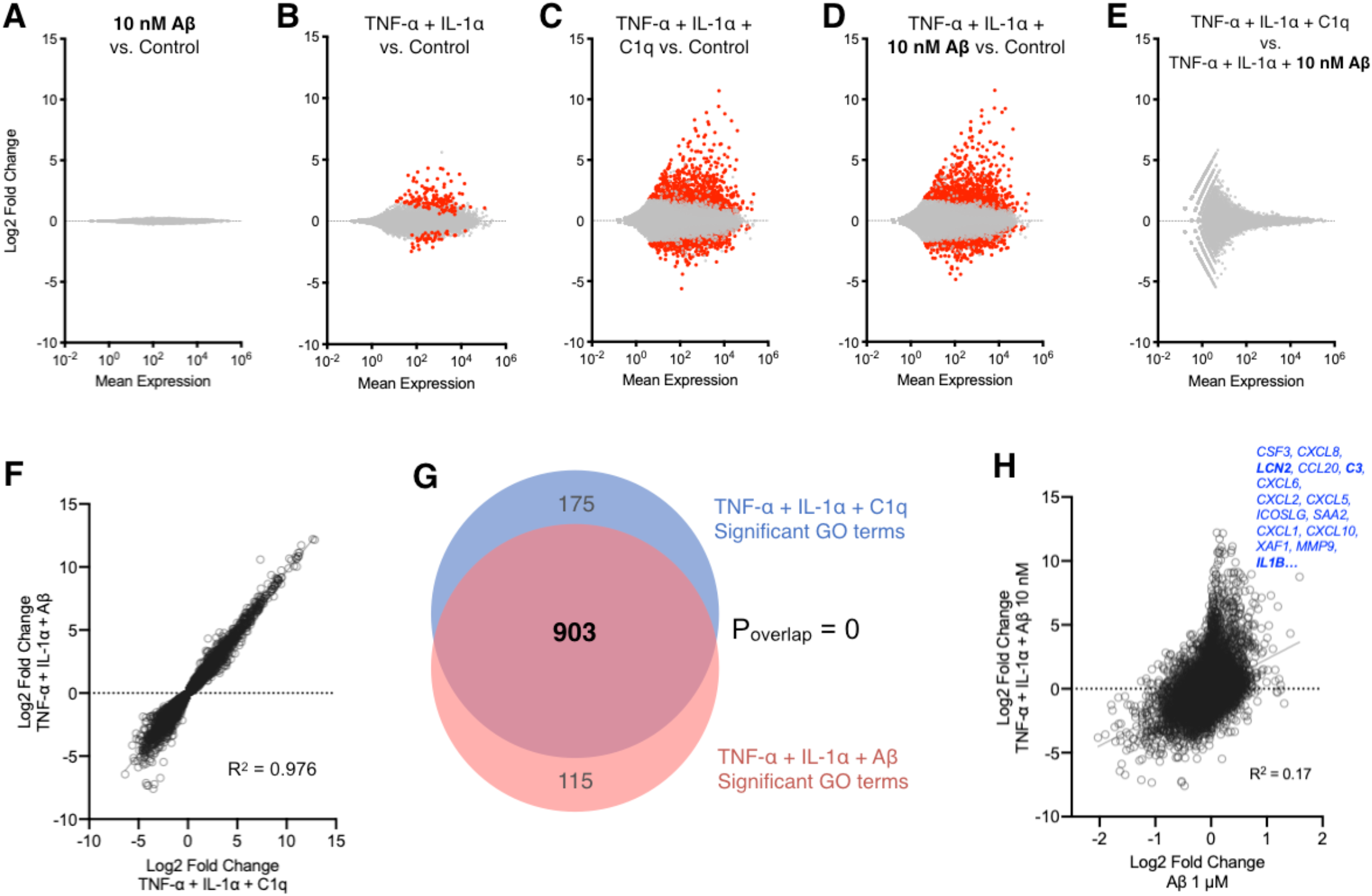
Similar transcriptional changes with cytokines and Aβ/cytokine cocktails. **(A-E)** MA plots showing Log2 fold change and mean gene expression in astrocytes treated with low-dose (10 nM) Aβ and/or combinations of TNF-α, IL-1α or C1q. Note no significant transcriptome changes with 10 nM Aβ alone and modest changes with TNF-α+IL-1α, but major and similar gene expression differences with both TNF-α+IL-1α+C1q and TNF-α+IL-1α+Aβ. Significantly increased/decreased genes in red (FDR<0.1). **(F)** Correlation between gene expression changes induced by cytokine cocktail vs. Aβ/cytokine cocktail. **(G)** Comparison of gene ontology (GO) terms significantly increased/decreased by cytokine-only and Aβ/cytokine cocktail. **(H)** Weak correlation between gene expression changes induced by Aβ/cytokine cocktail vs. high concentration (1 µM) Aβ alone; transcripts most different with Aβ/cytokine cocktail vs. Aβ alone in blue. All experiments performed in primary human astrocytes from three different donors.

### Aβ/cytokine-activated astrocytes are similar to disease-associated astrocytes

To determine specifically which biological pathways may be activated when Aβ synergizes with cytokines, we examined the genes that were differentially expressed in response to the Aβ/cytokine cocktail vs. TNF-α and IL-1α alone (**Figure 4A**). While this approach yielded fewer significant hits, gene ontology analysis of these transcripts pointed clearly to immune activation and cytokine signaling (**Figure 4B**).

**Figure 4.**
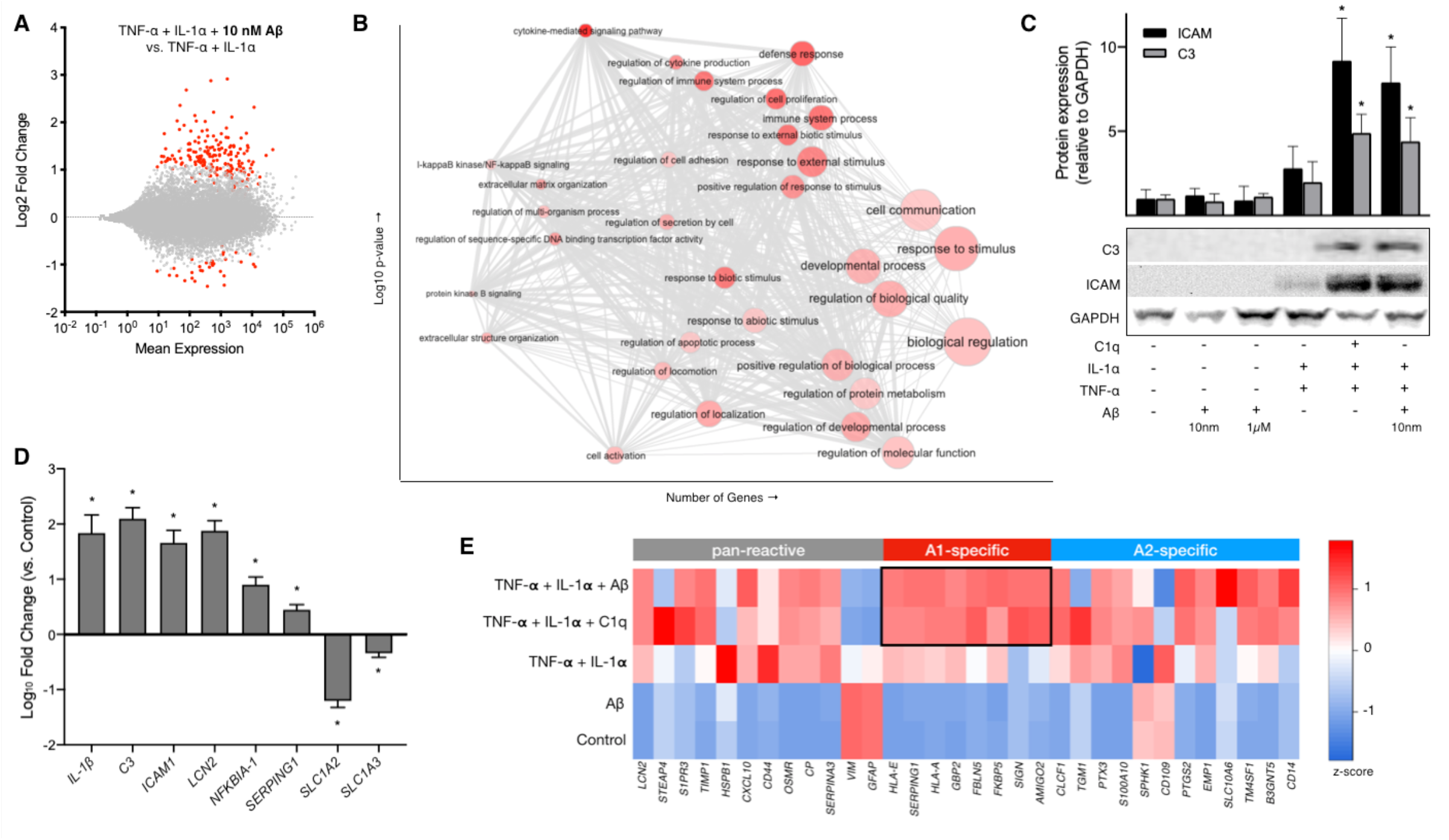
Upregulation of immune response and reactive astrocyte genes by Aβ/cytokine cocktail. **(A)** MA plot showing Log2 fold change and mean gene expression in astrocytes treated with TNF-α+IL-1α+Aβ vs. TNF-α+IL-1α alone (significantly increased/decreased genes in red, FDR<0.1; data on primary human astrocytes from three different donors, as in Figure 3). **(B)** Enhanced immune activation/cytokine signaling indicated by gene ontology analysis of genes differentially expressed when Aβ is added to TNF-α+IL-1α (based on significantly modulated transcripts in A). **(C-D)** Western blot and quantitative RT-PCR conifrmation of reactive astrocyte markers and innate immune mediators identified as increased/decreased with Aβ/cytokine cocktail treatment in RNA-seq data. Data collected using RNA and protein lysate from the same cells (one donor in triplicate) analyzed in Figure 1 and Supplementary Figure 1. *P<0.05 vs. control, unpaired two-tailed t-test. **(E)** Heat map showing relative expression of reactive astrocyte markers in response to Aβ and/or cytokines based on RNA-seq data from primary human astrocytes (one donor in triplicate as in Figure 1 and Supplementary Figure 1).

We next asked if these Aβ/cytokine-activated human astrocytes could be equivalent to the reactive, neurotoxic “A1” astrocytes induced by cytokine treatment of murine astrocytes. To test this possibility, we performed RNA-seq on Aβ/cytokine-treated astrocytes from one biological donor in triplicate (as in Figure 1), in order to reduce variability and increase resolution of key biological pathways. We found that these cells responded to Aβ/cytokine treatment with transcriptomic changes similar to those observed in our three-donor experiments (**Supplementary Figure 1**), and that gene expression differences in single vs. three-donor experiments were highly correlated (R^2^=0.84, p<0.000001), supporting this approach. We then confirmed changes in key genes that increased/decreased in both these and our three-donor data by both immunoblotting and RT-PCR, with a particular focus on mediators of innate immune activation and established markers of reactive astrocytes. We confirmed increased protein levels for two key reactive astrocyte markers (C3 and ICAM-1) in response to Aβ/cytokine treatment and the cytokine-only cocktail (**Figure 4C**). We also confirmed that Aβ/cytokine treatment caused significant increases in mRNA for: IL1B (IL-1β) and C3 (innate immune response mediators); nuclear factor κ B (NFκB, a major pro-inflammatory transcription factor); and *ICAM-1*, serpin G1 (*SERPING1*) and *LCN2*, all markers of reactive astrocytes [27, 40] (**Figure 4D**). Reduced transcripts confirmed by RT-PCR included excitatory amino acid transporters 1 and 2 (*SLC1A2*/*GLUT1* and *SLC1A3*/*GLUT2*), in line with our gene ontology analyses showing altered expression of synaptic function modulators. Interestingly, we noted that 1 µM Aβ did not affect the expression of IL-1β and C3 (**Figure 4C**), which is consistent with our initial RNA-seq data on this treatment. This observation further supports the idea that while Aβ may have effects on its own at high concentrations, it has an important biological function that may involve synergy with known pro-inflammatory cytokines at lower, physiologically relevant doses.

Finally, having confirmed that human astrocytes treated with the Aβ/cytokine cocktail expressed classic markers of astrocyte activation, we analyzed our RNA-seq data to determine if the treated astrocytes resembled “A1” reactive astrocytes that are neurotoxic, or “A2” activated astrocytes that are reportedly protective [27]. Both types of astrocyte are marked by activation of specific genes/proteins, as well as a suite of common response factors (“pan-reactive” astrocyte markers). As observed in previous mouse studies, we found that astrocytes treated with TNF-α+IL-1α+C1q increased expression of the full suite of A1 astrocyte markers. Moreover, in support of the idea that Aβ may contribute to astrocyte toxicity, we also found that Aβ/cytokine-treated astrocytes increased expression of this same set of markers (**Figure 4E**).

## DISCUSSION

Our studies extend on previous data [27, 28] by showing that a synergistic TNF-α+IL-1α+C1q cytokine cocktail induces immune activation in human astrocytes, and by demonstrating that Aβ may have an important, previously unrecognized role in neuroinflammatory signaling. This finding is highlighted by our observation that Aβ, at plausible physiological levels, can substitute for C1q in the activation of human astrocytes.

The idea that a potential “natural” function of Aβ contributes to AD pathology is new but gaining traction. Indeed, many earlier studies have focused on the pro-inflammatory, cellular effects of Aβ at high micromolar (aggregation-prone) concentrations. Here, we extend on those data with a whole-transcriptome (RNA-seq) analysis of biological pathways that may be involved in the response to high-dose Aβ. For example, we note that major transcriptional modules upregulated by Aβ in our data include numerous genes related to interferon response factors (IRFs), TNF proteins and receptors, and NF-κB signaling—many of which have been linked with Aβ before, but not in the same dataset. These and others’ data on the effects of high concentration Aβ may be relevant in the context of late-stage AD (i.e., when Aβ aggregates). However, many groups are beginning to reconsider the role of Aβ in AD through the lens of its potential/reported physiological functions [3], and the number of studies investigating the effects of Aβ at physiologically relevant concentrations is growing. These more recent investigations have shown that nano- and even picomolar concentrations of Aβ can modulate neuronal signaling and microglial activity [41, 42]. Our data are consistent with these recent reports, but they also provide new evidence for distinct biological effects of low concentration Aβ on astrocytes, which are increasingly implicated in brain aging and neurodegenerative diseases like AD [14, 28, 43]. In fact, taken together with recent evidence that innate immune activation may also drive Aβ production [44], our data could suggest a feed-forward model in which Aβ exacerbates immune activation in astrocytes and immune-activated astrocytes produce more Aβ. Our studies also raise a number of important questions:

First, to what extent do these cell culture experiments reflect *in vivo* cell physiology? Estimates of soluble Aβ peptide concentrations in the brain/cerebrospinal fluid are typically in the picomolar range [45, 46]. However, the existence of Aβ aggregates in senile plaques, coupled with biochemical studies demonstrating that concentrations of ~100 nM are required for Aβ aggregation [47], suggest that local Aβ concentrations significantly exceed the average brain Aβ concentration [48]. We note that we have used 10 nM levels of both cytokines and Aβ in our studies, and thus the synergistic pro-inflammatory effects of these peptides in cultured astrocytes are occurring at equivalent concentrations. We administered these treatments for 24 h, as this timepoint caused the greatest activation in preliminary studies, similar to what others have observed, but future studies could explore even lower concentrations and longer duration treatments that might further mimic *in vivo* physiology. The primary human astrocytes used in our study were from fetal sources, raising the possible concern that they may not fully reflect adult brain astrocytes. However, these primary cells are arguably: 1) more reflective of *in vivo* physiology than any related cell lines; and 2) a better model than aged and/or IPSC-derived astrocytes for investigating the *basic* biological effects of Aβ/cytokines, as their fundamental biological signaling pathways may be more intact than cells subjected to aging/reprogramming/other changes. Indeed, fetal CNS cells are typically used in cell culture experiments (e.g., rat embryonic hippocampal cultures) for these and other reasons. We cannot exclude the possibility that adult astrocytes would respond differently, but we are unaware of a specific mechanism that would produce this result. We also note that a recent report demonstrates that hIPSC-derived astrocytes from adult donors can be converted to neurotoxic, A1 astrocytes with the same TNF-α+IL-1α+C1q cocktail used in our studies [49], and all of the gene/transcript changes we confirmed in our data (Figure 4) follow the same pattern in RNA-seq data from that study. Interestingly, in both our data and this recent publication, some A1 astrocyte genes in reported in mice (e.g., *GFAP*) are not increased with cytokine treatment; this may be an important area for future study (i.e., determining specific differences between murine and human reactive astrocytes). Indeed, recent evidence suggests that AD-associated glial gene expression patterns may be markedly different in humans vs. mice [50] (although we note that all A1 astrocyte genes identified in mice that also have human orthologs were increased in cytokine-treated cells in our data).

Next, what species of Aβ are responsible for its pro-inflammatory effects? Our studies have employed nominally monomeric Aβ_1-42_, freshly prepared from dried HFIP-solubilized synthetic peptide. As we cannot control multimerization of the peptide during its 24 h incubation with cultured astrocytes, we cannot directly infer that the Aβ monomer is the pro-inflammatory species (although this could be more consistent with a role as a secreted cytokine-like molecule). We note that treatment with intentionally oligomerized Aβ using the media pre-incubation protocol originally described by Klein and colleagues [51] caused similar levels of immune marker induction (Figure 2B). Identification of the specific Aβ species that lead to immune activation will likely require additional studies that employ other versions of the Aβ peptide believed to have reduced (e.g., Aβ_1-40_, Aβ G37L) or increased (Aβ E22 delta) oligomerization capacity [52–54].

How could Aβ (or specific Aβ species) act in concert with cytokines to synergistically promote astrocyte activation? This is the salient question arising from our studies. Conceptually, Aβ could be acting through two different (but not mutually exclusive) types of mechanism: 1) direct interaction via an Aβ-specific receptor, activating an intracellular signaling cascade that converges with pathways stimulated by TNF-α, IL-1-α and C1q; or 2) by enhancing signaling downstream of these cytokines, perhaps by sensitizing the relevant receptors. There is a complex and sometimes contradictory literature on possible Aβ receptors [55, 56], and few of the associated studies have used astrocytes. Receptors and intracellular signaling pathways for the microglial cytokines we have used are much better understood, although their complexity confounds the ready identification of an obvious target for intervention. Thus, it may be necessary to conduct a large scale, unbiased genetic screen (e.g., a full-genome lentivirus-based CRISPR study) to determine the mechanism by which Aβ may act as a pro-inflammatory cytokine.

Finally, are the pro-inflammatory effects of Aβ/cytokines on astrocytes relevant to Alzheimer’s disease? Recent findings associating late onset AD (LOAD) risk alleles (e.g., in TREM2, CD33, ApoE, etc.) with microglial function [57, 58] have highlighted the likely role of neuroinflammation in AD. Astrogliosis is a common component of AD pathology, and the detection of putative neurotoxic “A1” astrocytes in aging and AD brains [27, 28] suggests a mechanism by which neuroinflammation could have a primary role in neuropathology (i.e., immune-activated glial cells causing neuronal death, rather than the other way around). Moreover, a specific population of pro-inflammatory “disease-associated astrocytes” has recently been detected in the context of both brain aging and AD [14], and we note that the most significantly upregulated genes in these disease-associated astrocytes (e.g., *GFAP, APOE, CST3*, etc.) are also significantly upregulated by the Aβ/cytokine cocktail used in our experiments. Thus, our studies suggest a model in which the accumulation of Aβ in AD could have a dual role in inducing neuroinflammation: accumulation of insoluble Aβ in senile plaques can lead to microglial immune activation, while synergy between soluble Aβ and microglial-secreted cytokines promotes the induction of neurotoxic “A1” astrocytes. Interestingly, deletion of the TNF-α receptor (*TNFR1*) has been reported to prevent the memory and neuronal dysfunction induced by injection of Aβ oligomers into mice [59], which is consistent with the hypothesis that the neurotoxicity of Aβ could in part stem from its synergistic pro-inflammatory interaction with TNF-α.

Collectively, our findings have potentially broader implications in that they could relate to the etiology of age-related (sporadic) AD. While there is a strong consensus that Aβ accumulation plays a central role in the initiation of AD pathology, the causes of this accumulation have not been established. If the Aβ peptide is indeed a natural, evolutionarily selected immune modulator, insults that provoke an innate immune response might be expected to induce increased production of Aβ. Numerous studies have associated bacteria or viruses with AD [60–62]. These studies have largely relied on analysis of postmortem brain samples, and thus cannot demonstrate causality. However, a recent study has shown that low level infection of a human cortical brain model with herpes simplex virus type 1 (HSV-1) results in gliosis accompanied by increased expression of both Aβ_1-42_ and TNF-α [63]. It is therefore tempting to speculate that microbial exposure in the brain could play a role in AD pathology via innate immune activation that upregulates Aβ, and that the age-related accumulation of Aβ (which eventually leads to aggregation/plaques) might be a consequence of these events.

## MATERIALS AND METHODS

### Astrocyte cell culture and treatments

Primary human astrocytes were used for all experiments. Cells from three different donors were purchased from different vendors (Lonza, lot number 0000647218; ScienCell, lot 22156; and Fisher/Gibco, lot 1948466). Vendors confirmed similar procedures for astrocyte isolation based on previously established protocols [64], in which the cerebral cortex was dissected, washed, minced, enzymatically digested to remove connective tissue and dissociated into astrocyte-specific medium in culture flasks and grown for several weeks, followed by isolation of adherent astrocytes by removing floating neurospheres (including via agitation), replating and expansion. All cells had typical astrocyte morphology and tested ~90% positive for GFAP (**Supplementary Figure 2**). We confirmed GFAP expression and also immuno-stained cells for fibroblast-specific protein 1 (FSP-1), but we observed minimal/no staining for this non-astrocyte marker in any cultures. Astrocytes were maintained in complete astrocyte growth medium (ScienCell) at 37°C and 5% CO_2_ in a humidified incubator. Cells were subcultured when confluent and grown to ~90% confluency for all experiments. All experiments were performed in serum-free media on passage 2-3 cells; treatments were applied by adding fresh medium containing pre-mixed reagents, and cells were harvested after 24 h incubation as previously described [34]. Lyophilized cytokines (TNF-α, IL-1α, C1q) were obtained from Sigma-Aldrich, resuspended in molecular biology grade water to make stock solutions, aliquoted and frozen for use in experiments (final experimental concentrations: 0.03 ng/µL [TNF-α], 0.30 ng/µL [IL-1α] and 4 ng/µL [C1q]) [27]. Aβ_1-42_ peptides (Anaspec and Abcam) were prepared following standard protocols [65] as dried HFIP films, then dissolved in fresh, anhydrous DMSO (Sigma) to make a ~5 mM solution, diluted in cold F12 media at target concentrations either for immediate use (monomers) or incubation overnight to produce oligomers when indicated, as previously described [66]. All experiments were repeated 3-5 times.

### RNA-seq and gene expression analyses

RNA-seq and gene expression analyses were performed using standard methods as previously described [34]. After the indicated treatments, astrocyte cultures were rinsed with DPBS and lysed in Trizol (Thermo), and RNA was recovered with an RNA-specific spin column kit (Direct-Zol, Zymo Research) including a DNase I treatment to remove genomic DNA. Poly(A)-selected libraries were generated using sera-mag magnetic oligo(dT) beads (Therma Scientific) and Illumina TruSeq kits. All libraries were sequenced on an Illumina NovaSeq 6000 platform to produce >40 M 150-bp paired-end fastq reads per sample. These reads were trimmed and filtered with the fastp program [67], then aligned to the hg38 *Homo sapiens* genome using the STAR aligner [68]. Gene/read counts were generated using HTseq-count [69], and differential gene expression was analyzed with Deseq2 [70]. Differentially expressed genes were then analyzed for gene ontology enrichment using both the GOrilla algorithm [71] and DAVID [72], and visualized using ReviGO [73]. To confirm RNA-seq results by RT-PCR, cDNA was generated from ~1 ug RNA per condition with random hexamers using Super Script IV First-Strand cDNA Synthesis Reaction (Invitrogen) according to manufacturer’s instructions, and a linearity curve was generated for all replicates. 5 ng cDNA was then used for PCR reactions (PerfeCTa SYBR Green FastMix Low ROX; Quanta Biosciences), which were quantified on an ABI 7900 instrument and normalized to GAPDH for analysis/presentation. Primer sequences were: IL-1β forward ATGCACCTGTACGATCACTG, reverse ACAAAGGACATGGAGAACACC; C3 forward AGGCCAAAGATCAACTCACC, reverse ATAGTGTTCTTGGCATCCTGAG; ICAM-1 forward CAATGTGCTATTCAAACTGCCC, reverse CAGCGTAGGGTAAGGTTCTTG; LCN2 forward TGAGCACCAACTACAACCAG, reverse AGAGATTTGGAGAAGCGGATG; NFKBIA-1 forward GTCTACACTTAGCCTCTATCCATG, reverse AGGTCAGGATTTTGCAGGTC; SERPING1 forward GTCCTCCTCAATGCTATCTACCTG, reverse GTTTGGTCAATGAAATGGGCCAC; SLC1A2 forward TCTTCCCTGAAAACCTTGTCC, reverse AGTCACAGTCTCGTTCAACAG; SLC1A3 forward CCATGTGCTTCGGTTTTGTG, reverse AATCAGGAAGAGAATACCCACG.

### Immunofluorescence staining

Astrocytes were grown and treated on glass chamber slides (Nunc Lab-Tek) for immunofluorescence staining and imaging as previously described [34]. After 24 h treatments, medium was removed and cells were washed in DPBS. Cells were then fixed in 4% paraformaldehyde, washed and permeabilized in 0.25% Triton-X, washed again and blocked in 3% fetal bovine serum/3% normal goat serum (Jackson ImmunoResearch). Primary antibody (CD44, Abcam; 1:500 dilution) was diluted in 5% normal goat serum and added to slide wells for overnight incubation at 4°C, after which slides were washed in DPBS with 0.1% tween and incubated with secondary antibody (Alexafluor 555, Invitrogen; 1:1000 dilution in 5% normal goat serum) for 1 h, followed by DAPI (Sigma-Aldrich; 1 µg/mL in DPBS) for 10 min at room temperature. Cells were washed again before mounting with Prolong mounting medium (Invitrogen) and then imaged on a Zeiss Axioskop fluorescence microscope at 40X magnification. Per cell fluorescence was measured in five images per slide well for each treatment and normalized to the mean of all conditions.

### Immunoblotting

Western blotting was performed on whole-cell lysates as previously described [34]. Semi-confluent astrocyte cultures were lysed in ice cold radio-immunoprecipitation assay lysis buffer with protease and phosphatase inhibitors (Roche), and 10 µg protein was separated by electrophoresis on 4-12% polyacrylamide gels followed by transfer to nitrocellulose membranes (all gels/reagents from Bio-Rad). Membranes were then blocked (5% milk in TBS-Tween buffer) and incubated for 18 h with primary antibodies: CD44 (Abcam, 1:1000 dilution); GFAP (ThermoFisher, 1:3000); ICAM-1 (Novus Biologicals, 1:2000); C3 (Novus, 1:2000); IL-1β (Novus, 1:1000). Proteins were then detected with horseradish peroxidase-conjugated secondary antibodies (Jackson ImmunoResearch) and ECL chemiluminescent substrate (Pierce) on a ChemiDoc imager (Bio-Rad), and signal (protein expression) was normalized to glyceraldehyde-3-phosphate dehydrogenase (GAPDH; Cell Signaling, 1:1000).

### Statistical analyses

All figures were prepared with GraphPad Prism software, which was also used to perform statistical tests indicated in figure legends. Differentially expressed genes shown in MA plots were detected using Deseq2 software as described above [70], and gene ontology overlap significance (hypergeometric probability) was calculated using the GeneOverlap program [74].

## Supplemental Data

**Supplemental Figure 1.**
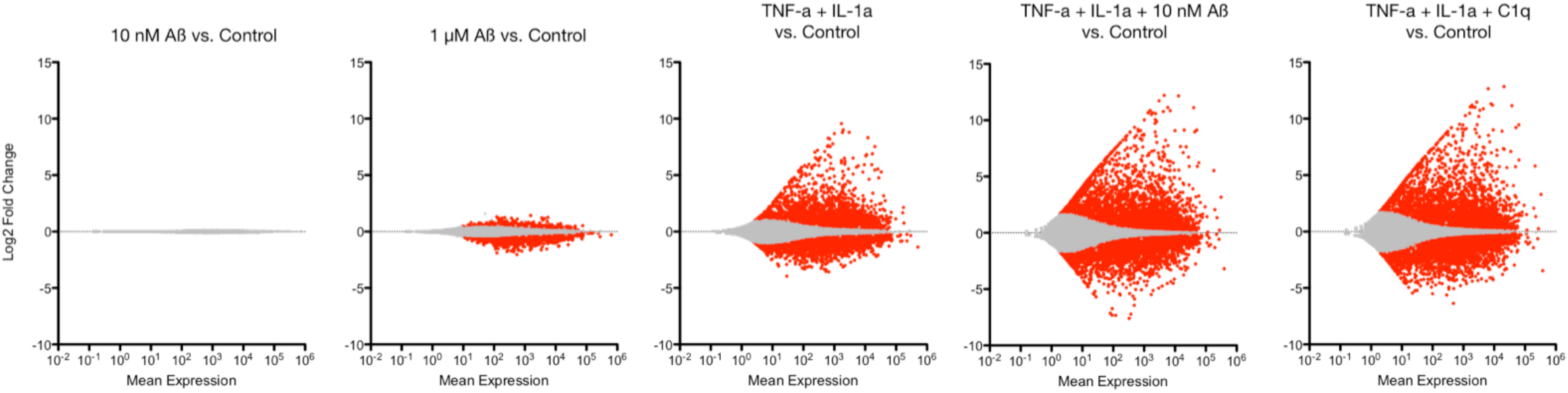
RNA-seq analyses of astrocytes treated with cytokine cocktail or Aβ/cytokines. MA plots showing Log2 fold change and mean gene expression in astrocytes treated with low concentration (10 nM) Aβ and/or combinations of TNF-α, IL-1α or C1q. Note no significant transcriptome changes with 10 nM Aβ alone and modest changes with TNF-α+IL-1α, but major and similar gene expression differences with both TNF-α+IL-1α+C1q and TNF-α+IL-1α+Aβ. Significantly increased/decreased genes in red (FDR<0.1). All experiments performed in triplicate with cells from one donor.

**Supplemental Figure 2.**
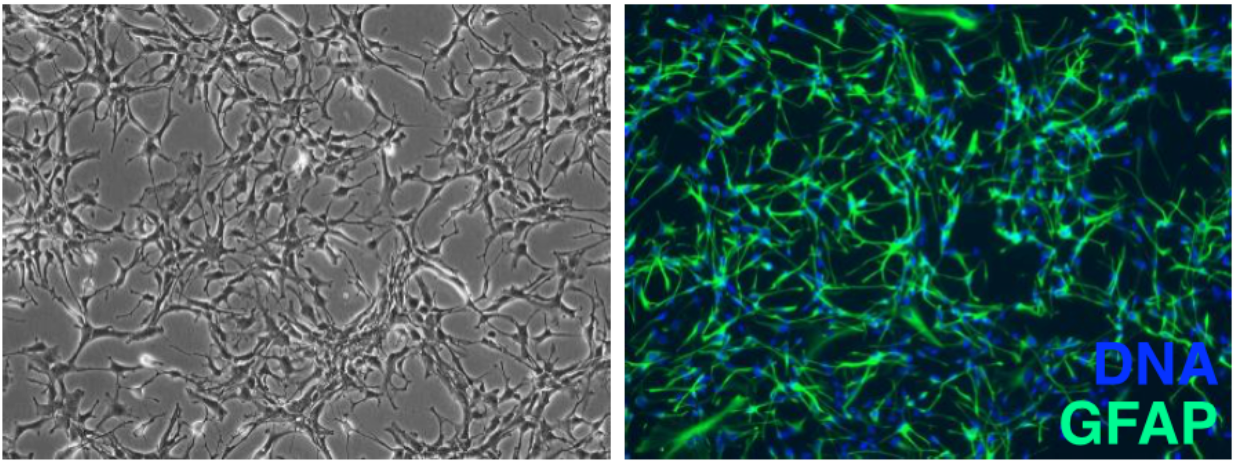
Representative images of primary astrocytes from commercial vendors. Phase contrast image (left) showing typical astrocyte morphology and immunofluorescence staining (right) showing positive staining for glial fibrillary acidic protein (GFAP) (example provided by ScienCell).

